# The mitochondrial unfolded protein response and metabolic reprogramming promote PASMC proliferation in response to sphingosine kinase-1/sphingosine-1-phosphate signaling

**DOI:** 10.1101/2025.06.10.658872

**Authors:** Angelia D. Lockett, Aaron Snow, Pontian Adogamhe, Justin Sysol, Manas Yadav, Tendo Mubuuke, Marta T. Gomes, Todd Cook, Amanda Fisher, Micheala A. Aldred, Roberto F. Machado

**Affiliations:** Division of Pulmonary, Critical Care, Sleep and Occupational Medicine, Indiana University School of Medicine, Indianapolis, IN, USA; Department of Medicine, University of Rochester Medical Center

**Author notes:** **Corresponding Author** Angelia D. Lockett, Indiana University Department of Medicine, Room C400B, Walther Hall, R3, 980 W. Walnut Street, Indianapolis, IN 46202. **Author Contributions** Conceptualization: Angelia D. Lockett, Micheala Aldred, Roberto F. Machado; Methodology: Angelia Lockett, Marta T. Gomes, Pontian Adogamhe, Roberto F. Machado; Data collection and analysis: Angelia D. Lockett, Aaron Snow, Pontian Adogamhe, Justin Sysol, Manas Yadav, Tendo Mubuuke, Marta T. Gomes; Writing-review and editing: Angelia D. Lockett, Micheala A. Aldred, Roberto F. Machado.

## Abstract

Proliferation and vasoconstriction of the intimal smooth muscle layer of the pulmonary artery are pathogenic characteristics of pulmonary arterial hypertension (PAH). Altered mitochondrial function, i.e. glycolysis, ROS generation and fission, are known potentiators of vascular remodeling. However, most current therapeutic interventions fail to effectively address the proliferation of the pulmonary artery smooth muscle cells (PASMCSs) lining the pulmonary vasculature and highlight the importance of identifying novel pathways to target for intervention. The Sphk1/S1P/S1P2 signaling axis is upregulated in PAH patients and is known to induce PASMC proliferation during hypoxia-mediated pulmonary hypertension (HPH). Interestingly, Sphk1 modulates mitochondrial function in that it regulates dynamics, cell growth and survival and in *C. elegans*, it induces activation of the unfolded protein response (UPR^mt^). We aimed to investigate if the Sphk1/S1P/S1P2 signaling axis promotes vascular remodeling in PAH via activation of the UPR^mt^. PASMCs isolated from IPAH patients were subjected to RNAseq analysis. The effect of Sphk1 or S1P was interrogated in hPASMC cell lines and the HPH model was used to assess the effect of the UPR^mt^ on PAH pathogenesis. RNAseq analysis revealed that pathways involved in mitochondrial respiration were among the top 20 most significantly regulated pathways. Furthermore, ATF-5, the transcription factor that mediates the UPR^mt^ was significantly upregulated. In hPASMCs, Sphk1/S1P lead to decreased respiration, increased glycolysis, fission, ROS and UPR^mt^ activation. Pharmacological inhibition of the UPR^mt^ mediator, mtHSP70, mitigated the Sphk1 induced increase in hPASMC proliferation. Furthermore, mtHSP70 inhibition was protective in hypoxia-mediated pulmonary hypertension (HPH) as we observed a decrease in right ventricular systolic pressure, right ventricular hypertrophy and vascular remodeling. These data suggest that the UPR^mt^ promotes vascular remodeling in PAH and may present a novel pathway to target for pharmaceutical intervention.

## Introduction

Pulmonary arterial hypertension (PAH) is an uncurable disease with a high mortality rate ranging from 18-55% over 3 years in patients with moderate to severe disease^1^. Death results from increased pulmonary vascular resistance (PVR) that eventually leads to right heart failure. Pulmonary arterial smooth muscle cells undergo changes in intracellular signaling that lead to a proliferative, apoptosis resistant phenotype that causes remodeling and occlusion of the pulmonary vasculature leading to increased PVR. The available therapies primarily address pulmonary vasoconstriction but have minor effects on alleviating other detrimental pathogenic characteristics of PAH, such as excessive pulmonary vascular remodeling.

We demonstrated that the Sphk1/S1P/S1PR2 signaling axis promotes vascular remodeling to mediate PAH development^2-4^. Sphingosine-1 Phosphate (S1P) is a bioactive sphingolipid that regulates cell proliferation, differentiation, motility and vascular tone and is generated by sphingosine kinase 1 (Sphk1) phosphorylation of sphingosine. Sphk1 expression is increased in the lungs and pulmonary artery smooth muscle cells (PASMCs) of PAH patients and in the lungs of rodent models of hypoxia mediated pulmonary hypertension (HPH). Consistent with this observation, S1P levels are also enhanced in the lungs of PAH patients and in the pulmonary arteries of rodents with HPH^3^. Metabolic reprogramming of pulmonary vascular cells also contributes to PAH via altered regulation of multiple mitochondrial processes, which leads to increased vascular cell proliferation and impaired vasorelaxation (for review see^5,6^).

Sphk1 has been shown to regulate mitochondrial function in Hela cells by promoting cleavage of mitofusin 2 (Mfn2)^7^. Cleavage of Mfn2 occurs during the process of mitochondrial fission. Interestingly, the calcium calmodulin domain of Sphk1 is thought to target Sphk1 to mitochondria to promote UPR^mt^ activation in the intestine and its kinase activity is required for UPR^mt^ activation in *C. elegans*^8^. The UPR^mt^ is a conserved transcriptional response that is regulated by nuclear-mitochondrial communication via shuttling of the activating transcription factor, ATF-5, between the cytoplasm and mitochondria. Perturbations in mitochondrial function such as oxidative phosphorylation defects, ROS and hypoxia lead to stabilization of ATF-5 translation in the cytoplasm which promotes its nuclear localization and initiation of the UPR^mt^ transcriptional program to restore mitochondrial homeostasis and promote cell survival ^4,9^. We hypothesized that the Sphk1/S1P signaling axis mediates metabolic reprogramming of PASMCs and promotes activation of the UPR^mt^ to promote vascular remodeling in PAH.

## Methods

### Reagents, Pharmacologic Inhibitors, and Antibodies

The following antibodies were purchased from Cell Signaling Biotechnology (Danvers, MA): HSP60 (12165s), ClpP (14181s), eIF2α (2103s), PCNA (13110s), p-Drp1 (3452s/3455s), Drp1 (5391s), mitofusin2 (11925s) anti-mouse-HRP (7076s), anti-rabbit-HRP (7074s), and Lamin-B2-HRP (24209s). MtHSP70 is from Invitrogen (Carlsbad, CA; MA3-028). ATF-5 (NBP2-67767) and S1PR2 (NBP2-26691) were purchased from Novus Biological (Centennial, CO). mtCO2 (ab110258) and p-eIF2α (ab32157) are from Abcam (Waltham, MA). β-Actin (A3854) and vinculin (CP74) were purchased from MilliporeSigma (Burlington, MA). DDK is from Origene Technologies (Rockville, MA; TA50011-100). Sphk1 was purchased from Proteintech (Rosemont, IL; 10670-1-AP). The Seahorse XF Glycolysis Stress Test Kit (103020-100), Seahorse XF Cell Mitochondrial Stress Test Kit (103015-100) and Seahorse XF base medium (102353-100) were purchased from Agilent Technologies (Santa Clara, CA). The Mitochondrial Extraction Kit is from BioVision Incorporated (Milipas, CA).

### Patient PASMCs and hPASMC Cell lines

Approval for the use of human lung cells was granted by the Indiana University Institutional Review Board. Deidentified human pulmonary artery smooth muscle cells isolated from the lungs of idiopathic pulmonary hypertension (IPAH, n=4), or control (n=4) patients were procured from the Pulmonary Hypertension Breakthrough Initiative (Indiana University, Indianapolis, IN). Primary hPASMC cell lines were purchased from Lonza (Walkersville, MA; CC-2581) or ATCC (Manassas, VA; PCS-100-023). Cells were cultured at 37°C in Smooth Muscle Cell Growth Medium (SmGm, Lonza, CC-3182) and were used between passage 4 to 8 for all studies.

### RNAseq Analysis

Total RNA was collected from IPAH-PASMCs and non-disease control PASMCs isolated from female individuals of European or African American descent. The Zymo RNA isolation kit was used (Zymo Research, Irvine, CA.; R1056). Three replicates were isolated per treatment group in two batches. Each batch contained patient and control cells from both ancestral backgrounds. Batch effect was corrected for in the RNAseq analysis. Total RNA was submitted to the Indiana University (IU) School of Medicine Center for Medical Genomics (Indianapolis, IN) for quality control, and RNA sequencing (Illumina). Sequence alignment, differential gene expression and gene set enrichment analysis (GSEA) were performed by Vugene (Kaunas, Lithuania). Briefly, the IU genomics core prepared the library using the KAPA mRNA Hyperprep Kit and ran the samples on a NovaSeq6000 for 100 base-paired end reads. FastQ files were provided to Vugene for analysis. Raw reads were aligned to the human GRCh38 version 44 genome using STAR aligner and FastQC and MultiQC sequence were used for quality reporting. For differential gene expression analysis, a mixed effects model with PatientID as a random effect and disease condition, age, sex, race and batch as fixed effects in the model. Normalized gene counts were used to fit the linear regression model using *R* package *edgeR* v3.42.4^10-12^. For GSEA gene symbols were ranked according to the sign of their differential expression fold-change and log-transformed p-value [−1×*sing*(*FC*)×*log*(*p*)−1×sign(FC)×log(p)], prior to using the GSEA software for analysis (UC San Diego, CA; Broad Institute, MA). Statistics: The model was adjusted for disease condition, race, age and batch. The negative binomial generalized log-linear model was fitted by running the *glmQLFit* function. Empirical Bayes statistics were estimated using the *glmQLFTest* function. The Benjamin-Hochberg (BH) method was used to correct for multiple testing. Genes with q-value < 0.05 were deemed significant.

### Animal Models of Pulmonary Hypertension and Hemodynamic Measurements

All experiments were approved by the Indiana University Institutional Animal Care and Use Committee (IACUC). For hypoxia-mediated pulmonary hypertension studies, male and female C57BL/6 mice (6-8wk of age) were exposed to room air (normoxia) or to 10% O_2_ (hypoxia) in a ventilated chamber for 4 weeks. A subset of both normoxia and hypoxia animals were administered Mkt-077 (10mg/kg, I.P.) every 2 days (except weekends) during the 4-week duration. Right ventricular systolic pressure (RVSP) was determined by right heart catheterization using a Millar pressure transducer catheter. Right ventricular hypertrophy (RVH) was determined by dividing the right ventricular weight by tibia length. Pulmonary vascular remodeling was assessed on hematoxylin and eosin-stained lung slices. The Aperio ImageScope software was used to examine a minimum of 10 microscopic fields per slide. As previously described, vascular remodeling was calculated using the following equation: (external vessel area-internal vessel area)/external vessel area^2,13^.

### S1P treatment and lentiviral transduction

hPASMCs were exposed to S1P (MilliporeSigma, Burlington, MA; S9666) as indicated. Lentiviral Sphk1-DDK and control vectors were designed, packaged and purchased from Vector Builder Inc (Chicago, IL). Human Sphk1 was cloned into a lentiviral plasmid that expressed hSphk1-DDK downstream of the EF1A promoter and EGFP downstream of the CMV promoter (VB010000-9298rtf). hPASMCs (100,000 cells/well) were seeded onto 6-well plates and allowed to attach overnight in a tissue culture incubator. Lentiviral particles were diluted in SmGm with sufficient viral titer to achieve MOI 20 and then added to cells. After 24h the lentiviral containing media was replaced with SmGm and left for 48h followed by downstream applications.

### Real-Time PCR

Total RNA was isolated from hPASMCs using the RNeasy Mini kit from Qiagen (Valencia, CA; #74104) and quantified with a Nanodrop 2000 spectrophotometer (ThermoFisher Scientific). Two micrograms of RNA were reverse transcribed with the High-Capacity cDNA Reverse Transcription Kit (ThermoFisher Scientific, #4368814). Expression of all mRNAs were monitored using gene-specific TaqMan primer assays with GAPDH expression used as an internal control (ThermoFisher Scientific). The ΔΔCt method was used to quantify relative changes in gene expression.

### Extracellular flus assay (Seahorse Assay)

Seahorse assays were performed according to the manufacturer’s protocol using the Agilent XF Pro M FluxPak (XFe96 cartridges, XF Pro Cell Culture plates and XF Calibrant; 103777-100). Transduced (MOI 20, 48h) hPASMCs were trypsinized and replated at 12,000 cells/well onto 96 well cell culture plates and incubated overnight to form a confluent monolayer. Oxygen consumption rate was assessed using the Seahorse XF Cell Mitochondrial Stress Test Kit. The cartridge plate was incubated overnight in XF calibrant in a 37°C incubator without CO_2._ On the day of the assay, Seahorse XF base medium was prepared with glutamine (2mM), sodium pyruvate (1mM) and glucose (10mM). The pH was adjusted to 7.4 and the medium was warmed to 37°C prior to washing cells 2 times. Cells were incubated in 200uL of XF base medium at 37°C for 1 hour in the absence of CO_2_ prior to performing the assay. Oligomycin (100uM), FCCP (carbonyl cyanide 4-(trifluoromethoxy) phenylhydrazone, 100mM), and rotenone/antimycin-A (50uM) were added to cartridge wells prior to loading the plate into the Agilent Seahorse XF Pro Analyzer. The final concentrations were as follows: 1.5 mM oligomycin, 1uM FCCP, and .5mM rotenone/antimycin-A. Extracellular acidification rate was measured using the Seahorse XF Glycolysis Stress Test Kit. The cartridge and cells were prepared as described above. On the day of the assay Seahorse XF base medium was prepared with 2mM glutamine and the pH was adjusted to 7.4. The cartridge wells were loaded with glucose (100mM), oligomycin (100uM), and 2-deoxyglucose (200mM). The final concentrations after injection were as follows: 10mM glucose, 1uM oligomycin, and 50 mM 2-deoxyglucose.

### Fluorescent Microscopy

hPASMCs (50,000 cells/well, MOI 20 48h) were seeded on glass coverslips in 12-well tissue culture plates and transduced as described above or treated with S1P. A 1mM Mitotracker Red (ThermoFisher; M7512) or 5mM MitoSox red (ThermoFisher; M36008) stock solution was prepared in DMSO and then diluted to 1uM in Hank’s Balanced Salt Solution. Culture medium was aspirated from hPASMCs and 250nM Mitotracker or 1µM MitoSox was added for 30 min in a 37°C tissue culture incubator. Cells were rinsed with cold PBS and then fixed with 4% paraformaldehyde solution prepared in PBS. The cells were washed 3 times in PBS and mounted onto coverslips using Prolong Gold antifade reagent with DAPI (ThermoFisher; P36941). Images were collected at 20x resolution using the Nikon Eclipse Ti2 fluorescent microscope (Melville, NY). Mean cellular fluorescence was measured using the Fiji Image J Software.

### Whole cell extraction, nuclear extraction, mitochondrial extraction and Western blotting

For analysis of whole cell extracts, cells were rinsed with cold PBS, collected in 1x cell lysis buffer (Cell Signaling; 9803) containing 1x protease/phosphatase inhibitor cocktail (ThermoFisher; 78442), incubated on ice for 15 min and centrifuged at 14,000 x g for 5 min to collect protein. For nuclear extractions, cells were rinsed with cold PBS and centrifuged at 500 x g for 5 min in 1mL of cold PBS. Cell pellets were gently resuspended in nuclear extraction buffer (10mM Tris-HCl pH 8.0, 140mM NaCl, 1.5mM MgCl_2_, 0.5% NP-40 (v/v), 1x protease/phosphatase inhibitor cocktail) followed by incubation on ice for 5 min. The supernatant containing cytoplasmic proteins was transferred to a new Eppendorf tube. The nuclear pellet was resuspended in cell lysis buffer and incubated on ice for 30 min followed by centrifugation at 14,000 x g for 5 min. in a separate set of experiments, cells were rinsed with cold PBS followed by isolation of mitochondria using the Mitochondrial Isolation Kit according to the manufacturer’s protocol (BioVision). Proteins were resolved by SDS-PAGE and transferred to PVDF membranes. Western Blotting was performed by incubating with primary antibodies (1:1000, overnight, 4° Celsius). HRP conjugated secondary antibodies were used at 1:5000 for 1h at room temperature.

### Cell Proliferation

Following the completion of the transduction procedure as described above, Mkt-077 (1uM) was added to hPASMCs. Live cell images were collected at time 0 and at 4hr time intervals over a 48h period using the IncuCyte Live-Cell Analysis System (Sartorius, Bohemia, NY). Cell proliferation was assessed per the manufacturer’s instructions. Briefly, IncuCyte uses a series of masks to track nuclei number, percentage confluence, and GFP signal. Increasing nuclei count, and GFP intensity was used as a measure of cell proliferation.

### Mouse PASMC Isolation

mPASMCs were isolated using one lung per treatment group. The Jove PASMC isolation protocol was used as described with minor modifications^14^. Briefly, M199 media (complete and serum free), pulmonary artery (PA) agarose containing iron particles, lung agarose and collagenase were prepared as described. PA agarose was slowly perfused through the right ventricle to the pulmonary artery until the lung appeared gray. Lungs were inflated with lung agarose prior to removing the heart-lung bloc and solidifying in ice cold PBS. Lungs were minced (1 mouse per treatment group) in 1mL of ice-cold PBS in a tissue culture hood. Minced lungs were transferred to a 15mL conical tube placed in a magnet and washed with ice cold PBS. Tubes were removed from the magnet and iron pellets containing lung tissue were resuspended in collagenase medium and incubated at 37°C in a CO_2_ incubator for 1h. The tissue was disrupted by passing through a 15 G blunt needle attached to a 10mL syringe 5 times. The tissue-iron slurry was next passed through an 18 G blunt needle 5 times to completely disrupt visible tissue chunks. The tissue-iron slurry was transferred to a 15mL conical tube placed in a magnet and washed with complete M199 medium. The tube was removed from the wall and a compact tissue-iron pellet was resuspended in complete M199, transferred to a 35mM tissue culture dish (plate 0) and incubated overnight. The following day, the supernatant was transferred to a 15mL conical on a magnet, washed with complete medium and transferred to a new 35mM tissue culture dish (plate 1) and allowed to migrate from pulmonary artery tissue and proliferate until confluent (∼ 4 days). Cells were expanded and used between passage 1-3.

### Statistical Analysis

Statistical analysis of experimental data was performed using GraphPad Prism 5.1 (GraphPad Software, Inc., La Jolla, CA). Results are expressed as mean ± SEM from at least three experiments. Student *t* test and analysis of variance were used to compare two and three groups, respectively. *P* less than 0.05 was considered statistically significant.

## Results

### 1. Genes that regulate mitochondrial ROS and the UPR^mt^ are differentially expressed in PASMCs isolated from IPAH patients

We performed RNAseq analysis on PASMCs isolated from idiopathic pulmonary arterial hypertension (IPAH) patients and non-disease controls. Three pathways involved in aerobic respiration were among the top 20 most downregulated pathways in IPAH-PASMCs (Fig. 1a). Multiple genes, transcribed from both the nuclear and mitochondrial genome, which are essential to function of complex I of the electron complex train were significantly downregulated. Additionally, ATF-5, which transactivates genes that promote the UPR^mt^ was significantly upregulated (Fig. 1b). Although moderate, S1PR2 expression was also significantly increased. Our group previously demonstrated that S1PR2 is increased in the lungs of IPAH patients and that the Sphk1/S1P/S1PR2 signaling axis promotes vascular remodeling in PAH^3^. These data, together with our RNAseq observations, suggest that congruent with upregulation of the Sphk1/S1P/S1PR2 pathway, multiple mitochondrial processes are also altered in PAH patients.

**Figure 1.**
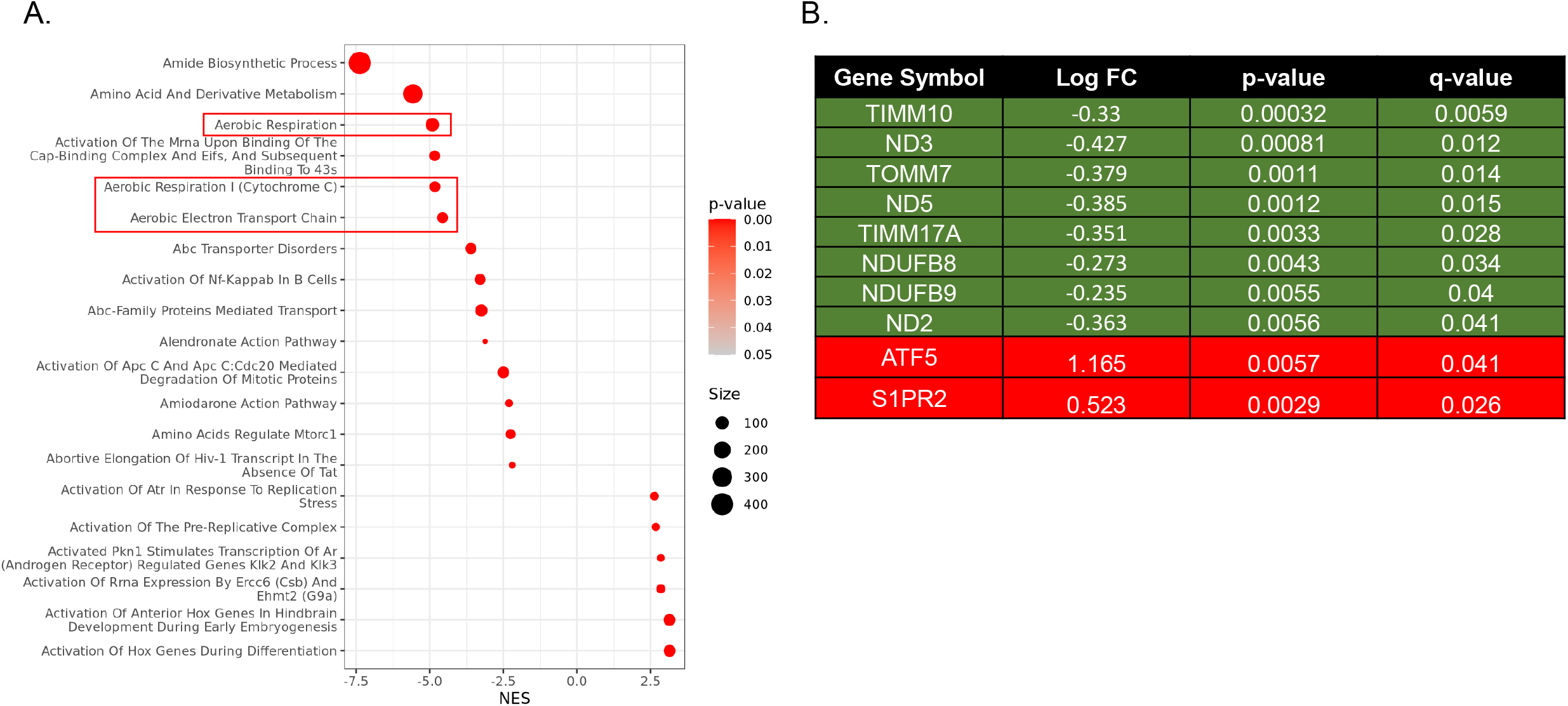
Normalized enrichment scores of the top 20 most significant pathways. Pulmonary artery smooth muscle cells isolated from the lungs of control and idiopathic pulmonary arterial hypertension (IPAH) patients were subjected to RNAseq analysis followed by gene set enrichment analysis. Red boxes indicate enriched mitochondrial respiration pathways (A). The table (B) highlights differential expression of genes within complex I of the electron transport chain, the UPR^mt^ pathway and the Sphk1/S1P/S1PR2 signaling axis. -NES/Down=decreased in IPAH; +NES/Up=increased in IPAH.

### 2. The Sphk1/S1P/S1P2 signaling axis modulates mitochondrial function in hPASMCs

HIF1α is a well-established mediator of PAH that is known to promote a metabolic switch in the pulmonary vasculature from mitochondrial respiration to glycolysis^15,16^. This, in turn, leads to PASMC proliferation and vascular remodeling. In multiple cancer cell lines and in liver cells, Sphk1 increases HIF1α expression to promote cell growth and survival^17,18^. Based on these studies and our RNAseq findings that both mitochondrial processes and S1PR2 are differentially expressed in IPAH-PASMCs, we assessed the effect of the Sphk1/S1P/S1PR2 pathway on HIF1α expression and mitochondrial function in hPASMCs. S1P treatment of hPASMCs promoted increased HIF1α gene expression and nuclear localization that was S1PR2 dependent (Fig 2a,b). Extracellular flux analysis (Seahorse assay) was performed on cells that were transduced with lentivirus packaged with either a control plasmid (Con) or Sphk1 expressing plasmid (Sphk1). Overexpression of Sphk1 lead to decreased oxygen consumption rate and increased glycolysis (Fig. 2c,d). In PAH, impaired respiration leads to ROS generation which increases vasoconstriction. Hence, we assessed the effect of Sphk1 on mitochondrial ROS production using mitoSox staining and found that Sphk1 lead to increased ROS generation (Fig. 2e). In addition to promoting glycolysis, HIF1α promotes PASMC proliferation in PAH by inducing mitochondrial fragmentation^19^. Hence, the effect of S1P on mitochondrial fission was assessed by exposing hPASMCs to S1P and measuring the activation state of Drp1 and the expression level of Mfn2. S1P lead to increased Drp1 phosphorylation concomitant with decreased Mfn2 expression, which is indicative of increased mitochondrial fragmentation which we observed using Mitotracker red staining (Fig 2f-h). Together, these data indicate that the Sphk1/S1P/S1P2 signaling axis regulates mitochondrial function in PASMCs via modulation of multiple mitochondrial processes.

**Figure 2.**
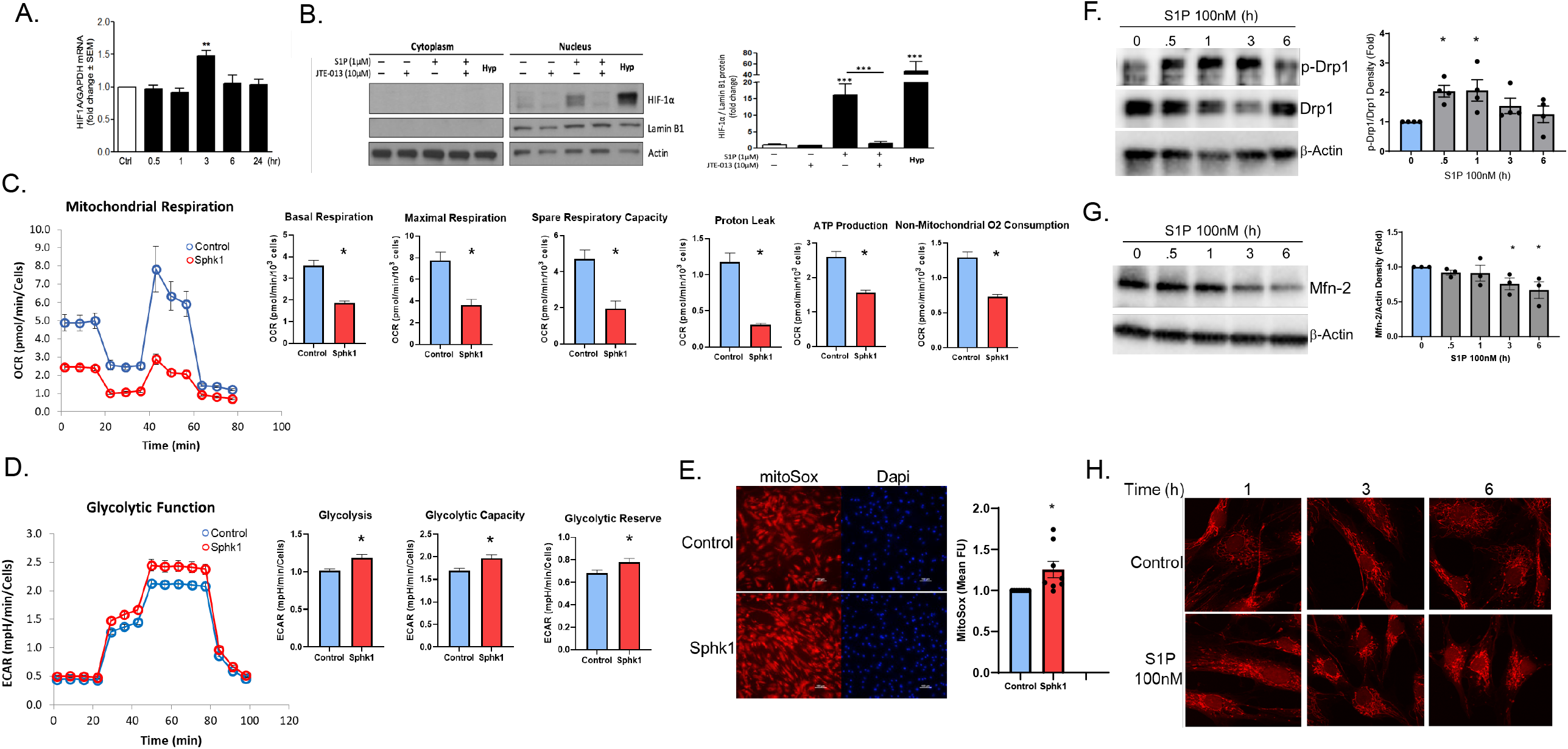
Sphk1/S1P promotes mitochondrial dysfunction in hPASMCs. hPASMCs were treated with S1P or transduced with Lenti-Sphk1 (MOI20, 48h). Real-time PCR (A) and Western blotting (B) demonstrate increased HIF1α expression and nuclear localization. Seahorse analysis for oxygen consumption rate (OCR) and extracellular acidification rate (ECAR), show that Sphk1 overexpression promoted decreased mitochondrial respiration (C) and increased glycolysis (D). Immunofluorescence for MitoSox Red (E) show that Sphk1 increased ROS. Western blotting (F-G) and immunofluorescence for Mitotracker Red (H) demonstrate that S1P promoted increased fission. Results are expressed as mean ± SEM versus control. p values are as indicated or ^*^,^**^,^***^p≤.05.

### 3. The Sphk1/S1P signaling promotes ATF-5 nuclear localization and UPR^mt^ activation

ATF-5 induced activation of the UPR^mt^ is involved in tumorigenesis of multiple types of cancer^4,20,21^. Both hypoxia and ROS, which are central features of metastasis, promote ATF-5 activation^4,22^. Phosphorylation of the translation initiation factor eIF2α, promotes ATF-5 cytosolic translation allowing for nuclear localization and transactivation in cancer cells. Given the importance of ATF-5 in cancer, and the synthetic phenotype of pulmonary vascular cells in PAH, we assessed the expression level of UPR^mt^ mediators in IPAH-PASMCs. Phosphorylated eIF2α trended upward, and total eIF2α was significantly increased in IPAH-PASMCs (Fig. 3a) suggesting adaptation to stress. Regarding UPR^mt^ proteins that localize to the mitochondria, while there was a trend toward increased expression, only HSP60 was significantly increased (Fig. 3). However, since both the ATF-5 and S1PR2 genes were differentially expressed in our IPAH-PASMC RNAseq data set, we used hPASMC cell lines to investigate the effect of Sphk1/S1P signaling on UPR^mt^ activation. Similar to our observations in patient PASMCs, both p-eIF2α and total eIF2α expression were increase in response to Sphk1 overexpression (Fig. 4a). Furthermore, Sphk1, which catalyzes the conversion of sphingosine to S1P, also lead to increased nuclear localization of ATF-5 (Fig. 4b). S1P lead to increased nuclear localization of ATF-5 at both acute and extended (24h) time points (Fig. 4c,d). 20 MOI was sufficient to overexpress Sphk1 as determined by Western blotting for both Sphk1 and its DDK-tag (supplementary Fig. 1).

**Figure 3.**
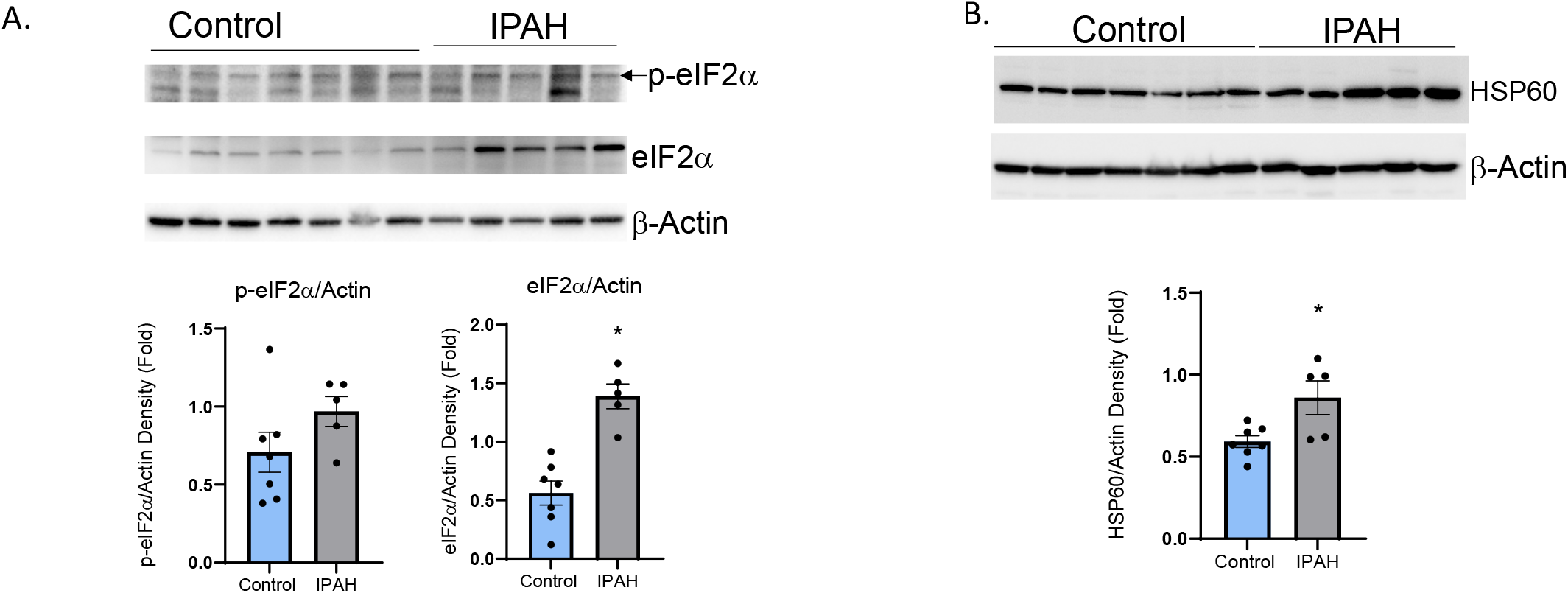
Expression of the UPR^mt^ mediator, HSP60, is increased in IPAH-PASMCs. Whole cell extracts were collected from PASMCs isolated from the lungs of control and IPAH patients followed by measuring the expression of p-eIF2α, eIF2α (A) or HSP60 (B). Western blot and densitometry demonstrate elevated protein levels of eIF2α and HSP60and a trend toward increased p-eIF2α. Results are expressed as mean ± SEM versus control. p values are as indicated or ^*^p≤.05.

**Figure 4.**
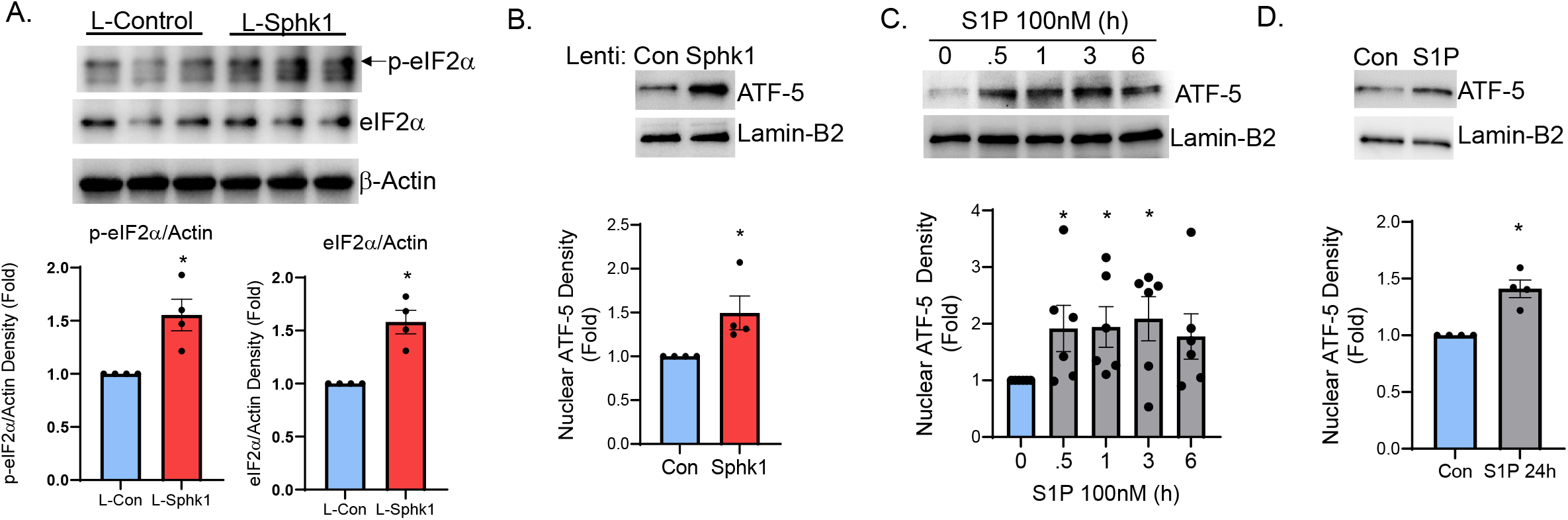
ATF-5 localizes to the nucleus in response to Sphk1/S1P stimulation. hPASMCs were transduced with Lenti-Sphk1 (MOI20, 48h) or exposed to S1P (.5-24h). Western blotting and densitometry of whole cell extracts demonstrate increased expression and phosphorylation of eIF2α (A). Western blotting and densitometry of nuclear fractions demonstrate ATF-5 translocation to the nucleus in response to Sphk1 overexpression (B) and S1P treatment (C,D). ^*^p≤.05 versus time 0 or control.

When the UPR^mt^ transcriptional program is activated, ATF-5 induces the expression of chaperones, such as mtHSP70 and HSP60 as well as proteases such as ClpP, that regulate mitochondrial proteostasis. These proteins translocate to damaged mitochondria to restore mitochondrial function and to promote cell survival^23^. Overexpression of Sphk1 lead to increased expression of mtHSP70, HSP60 and ClpP (Fig. 5a-c). Treatment with S1P confirmed the effect of Sphk1 on UPR^mt^ activation as it also enhanced expression of the UPR^mt^ mediators (Fig. 5e-g) and at time points that were consistent with the timing of ATF-5 nuclear localization (Fig. 4). Concomitant with UPR^mt^ activation, S1P and Sphk1 stimulated hPASMC proliferation as indicated by increased PCNA expression (Fig. 5d,h). Together, these observations indicate that Sphk1/S1P promotes both hPASMC proliferation and upregulation of the UPR^mt^ mediators. Hence, we next determined if UPR^mt^ activation promotes hPASMC proliferation.

**Figure 5.**
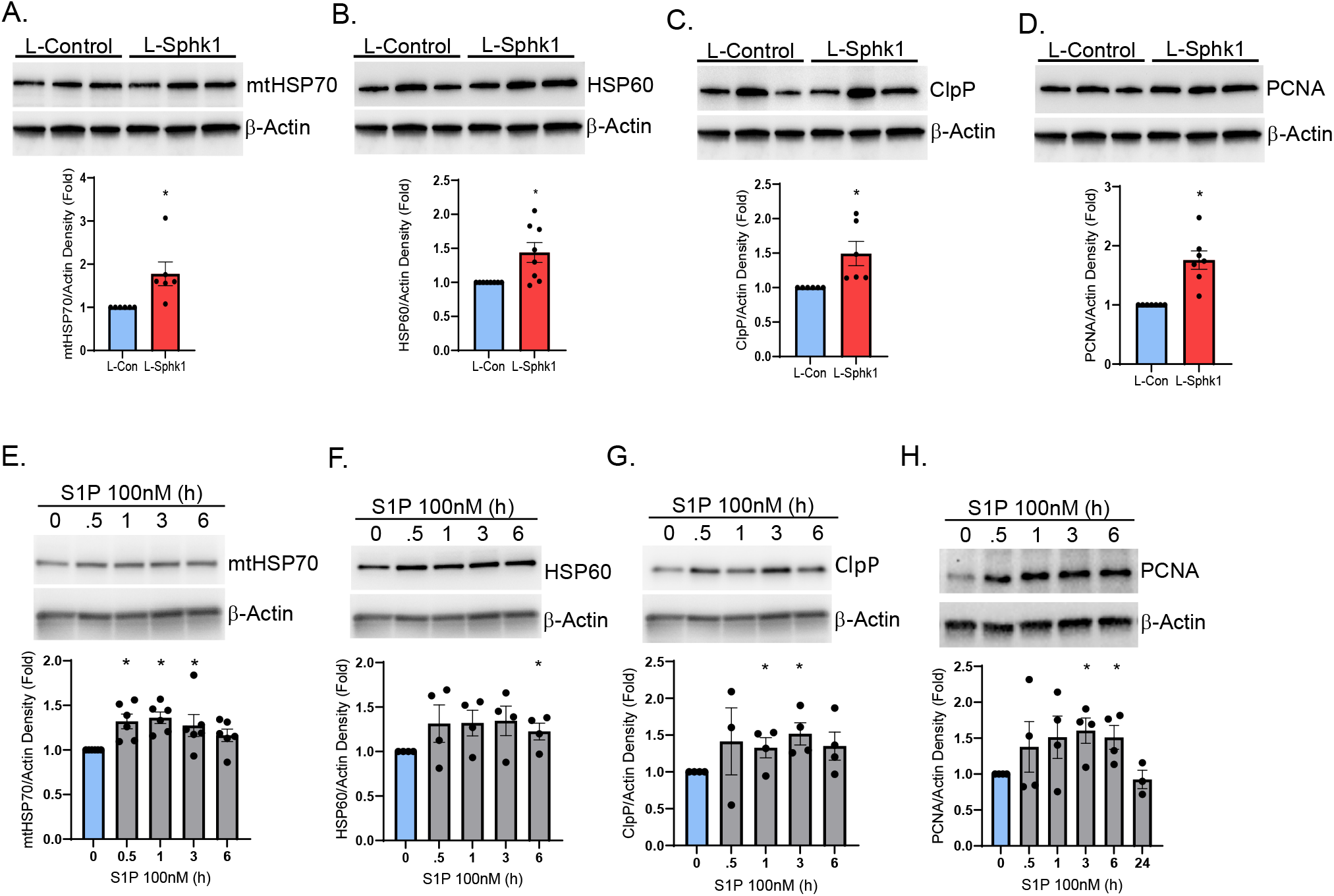
The expression and mitochondrial localization of UPR^mt^ is increased by Sphk1/S1P stimulation. hPASMCs were transduced with Lenti-Sphk1 (MOI20, 48h) or exposed to S1P. Western blotting and densitometry of whole cell extracts demonstrate that Sphk1 (A-C) and S1P (E-G) lead to increased expression of UPR^mt^ mediators and promoted proliferation (D,H). Results are expressed as mean ± SEM. ^*^p≤.05 versus time 0 or control.

### 4. Inhibition of the UPR^mt^ is protective against hPASMC proliferation

mtHSP70, also known as mortalin, is a HSP70 family member that primarily localizes to the mitochondria and can be specifically inhibited using the pharmacological agent, Mkt-077 which is an allosteric inhibitor that crosses the mitochondrial membrane. We investigated the effect of Mkt-077 on Sphk1 stimulated hPASMC proliferation by overexpressing Sphk1 for 48h followed by treatment with Mkt-077. hPASMC proliferation was assessed by Western blotting 24h after Mkt-077 addition or by live cell imaging immediately after adding Mkt-077, for up to 48h. Sphk1 lead to increased PCNA expression that was mitigated by mtHSP70 inhibition (Fig. 6a). The effect of UPR^mt^ on proliferation was confirmed by measuring fluorescence of live cells which demonstrates that inhibition of mtHSP70 attenuated Sphk1 induced hPASMC proliferation (Fig. 6c). To further investigate the effect of the UPR^mt^ on cell growth, ATF-5 and S1PR2 were knocked-down using siRNA. The effectiveness of the depletion was determined by immunoblotting for ATF-5 and S1PR2, both of which were significantly reduced (Fig. 6b). Both ATF-5 and S1PR2 depletion led to reduced baseline expression of PCNA (Fig.6b). These data indicate that Sphk1/S1P induced activation of the UPR^mt^ pathway promotes hPASMC proliferation; hence we next determined the involvement of the UPR^mt^ in mediating vascular remodeling *in vivo*.

**Figure 6.**
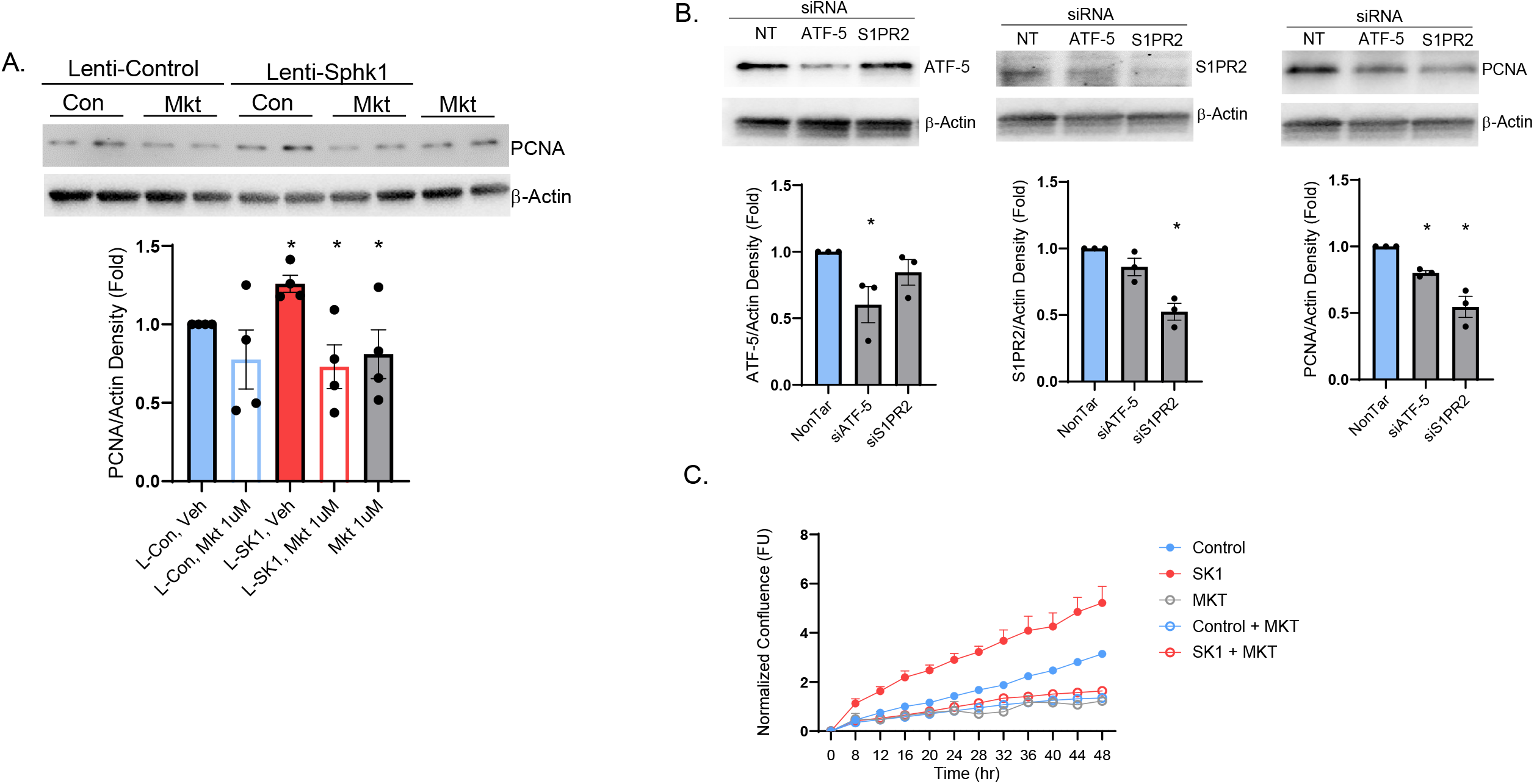
Inhibition of mtHSP70 mitigates Sphk1 induced hPASMC proliferation. Cells were transduced with Lenti-Sphk1 (MOI20, 48h) followed by treatment with Mkt-077 (1µM, 24-48h) or were transfected with siRNA for 72h. Western blotting and densitometry of whole cell extracts demonstrate that Sphk1 lead to increased expression of PCNA and that Mkt-077 mitigates the Sphk1 effect on PCNA expression (A). Western blotting and densitometry demonstrate that knock-down of ATF-5 or S1PR2 lead to decreased PCNA expression (B.) IncuCyte Live Cell Imaging analysis demonstrate that Mkt-077 prevented Sphk1 induced increase in proliferation over a 48h time period (C). Results are expressed as mean ± SEM. ^*^p≤.05 versus control.

### 5. Inhibition of the UPR^mt^ is protective against hypoxia-mediated pulmonary hypertension (HPH) development

To determine if UPR^mt^ activation occurs *in vivo*, C57BL/6 mice were exposed to room air, or 10% hypoxia for 4 weeks followed by isolation of PASMCs from mouse lungs. Compared to normoxic mPASMCs, we observed a significant increase in mtHSP70 expression (Fig. 7a). Therefore, we assessed whether inhibition of mtHSP70 could prevent HPH. C57BL/6 mice were exposed to room air or 10% hypoxia only, for 4 weeks or injected I.P. with Mkt-077 every 2 days during the 4-week exposure. Compared to normoxia-exposed control mice, hypoxia-exposed mice had elevated right ventricular systolic pressure (RVSP), right ventricular hypertrophy (RVP, RV/tibia length) and pulmonary vascular remodeling (Fig. 7b-f). Inhibition of the UPR^mt^ by treatment with Mkt-077 mitigated the increase in RVSP, RVP and vascular remodeling (Fig. 7b-f).

**Figure. 7.**
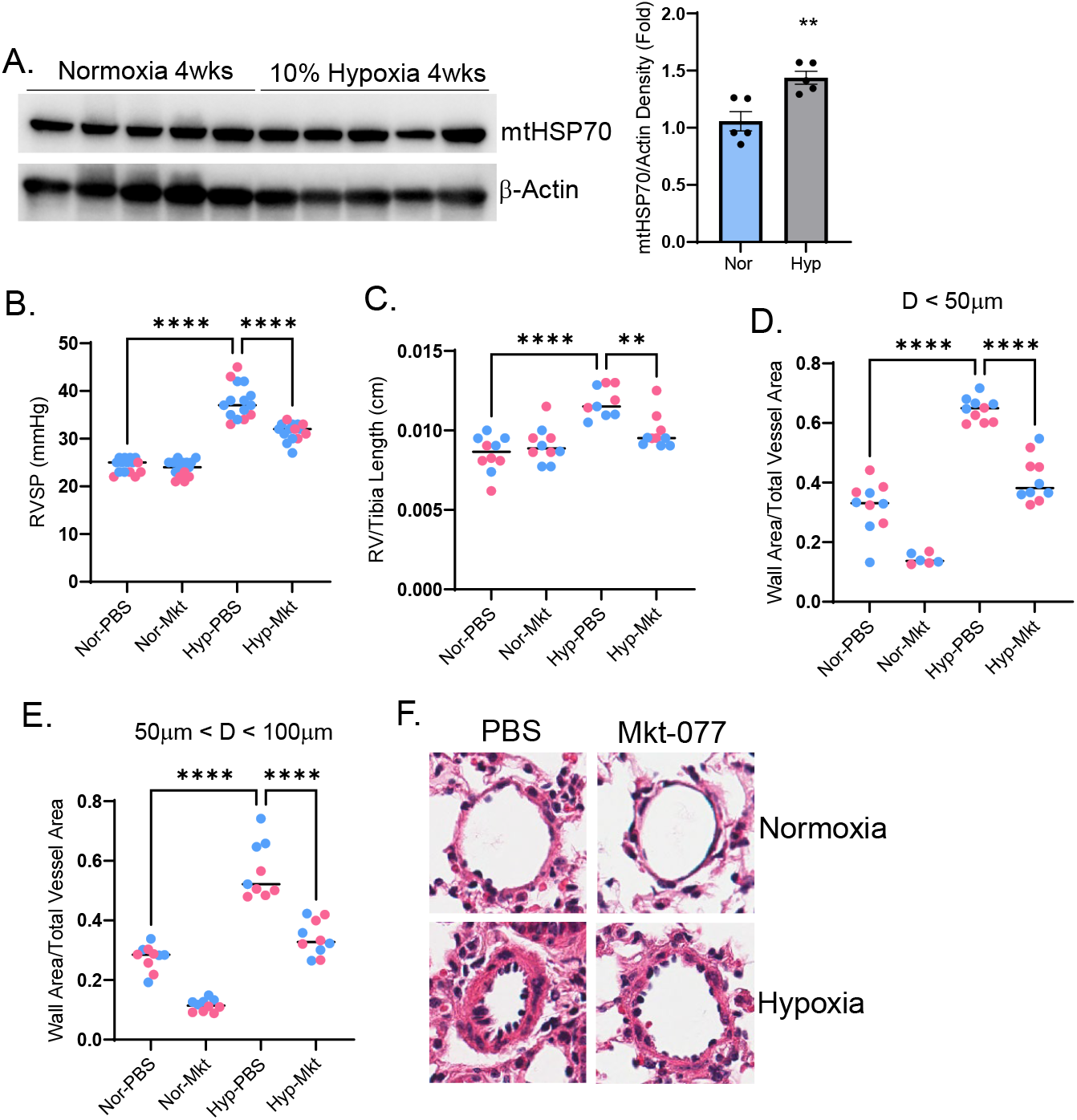
Inhibition of mtHSP70 blocks proliferation and mitigates hypoxia-mediated PH. (A) C57BL/6 mice were exposed to room air, or 10% hypoxia for 4 weeks followed by isolation of mouse PASMCs. Hypoxia promoted mtHSP70 expression as shown by Western blot and densitometry. (B-F) C57BL/6 mice were exposed to room air or 10% hypoxia for 4 weeks with or without Mkt-077 (I.P, 10mg/kg every 2 days). Hypoxia increased right ventricular systolic pressure (B), RV thickness (C) and vascular remodeling (D-F); all parameters were decreased by Mkt-077 administration. Results are expressed as mean ± SEM.^**^p≤.006, ^***^p≤.003,^****^p≤.0001.

**Figure 8.**
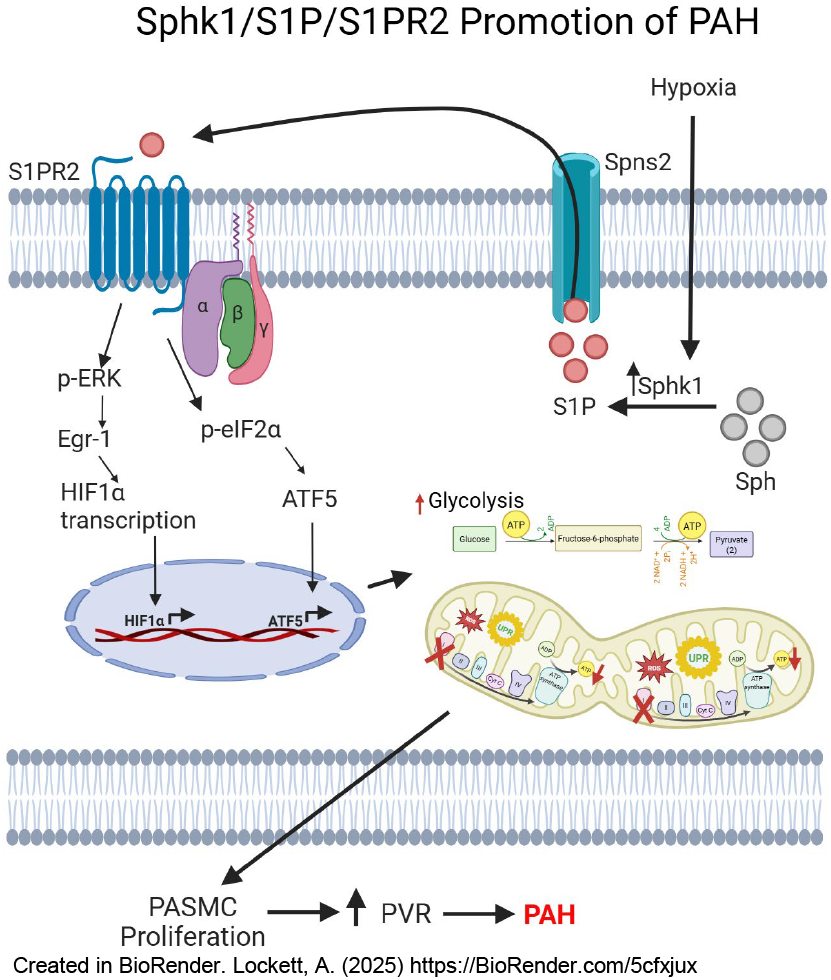
Schematic depicting Sphk1/S1P/S1PR2 modulation of mitochondrial function to promote vascular remodeling and PAH. In PASMCs hypoxia leads to upregulation of Sphk1, which catalyzes the conversion of sphingosine (sph) to sphingosine-1-phosphate (S1P). S1P is transported to the extracellular environment by the Spns2 transporter allowing S1P to engage its receptor, S1PR2, to stimulate intracellular signaling events. The ERK pathway is activated to induce Egr-1 translocation to the nucleus to transactivate HIF1α. HIF1α leads to decreased respiration and increased glycolysis and mitochondrial fragmentation to promote PASMC proliferation. Engagement of S1PR2 also leads to activation of eIF2α which induces ATF-5 nuclear localization and activation of a transcriptional program that promotes the UPR^mt^ and PASMC proliferation. During S1P signaling mitochondrial ROS is generated, which signals mitochondrial stress, a of stimulant of UPR^mt^ activation. The Sphk1/S1P/S1PR2 signaling axis leads to activation of two transcription programs via, HIF1α and ATF-5, to promote PASMC proliferation which leads to increased pulmonary vascular resistance to promote the PAH pathogenesis.

## Discussion

Herein we report that Sphk1/S1P/S1PR2 exerts regulatory effects on mitochondrial function via modulation of multiple mitochondrial processes: bioenergetics, dynamics and the stress response. In PAH patients, the Warburg effect, a glycolytic shift in energy production from aerobic respiration to glycolysis at normal oxygen concentrations, is associated with a highly proliferative PASMC phenotype^24,25^. Increased Sphk1 expression and signaling promotes glycolysis and oncogenesis in multiple types of cancer^26^. These functions of Sphk1 have been linked to the ability of Sphk1 to increase HIF1α^27^, a transcription factor that enhances metabolic reprogramming and cell migration and proliferation in PAH. We observed that S1P promoted nuclear localization of HIF1α which was dependent on S1PR2. We have previously shown that the Sphk1/S1P/S1PR2 pathway is upregulated in PAH patients and that it promotes abnormal PASMC growth and vascular remodeling in animal models of PH^3^. Our observations herein demonstrate that Sphk1/S1P induced metabolic reprogramming of PASMCs in that ATP generated via aerobic respiration is impaired concomitant with an increase in glycolysis. Consistent with the decrease in aerobic respiration, Sphk1 lead to increased mitochondrial reactive oxygen species (ROS) generation. Increased ROS in PAH promotes a pulmonary vascular environment that is conducive to hyper-proliferation, vascular occlusion, vasoconstriction and increased pulmonary arterial pressure^28-30^. Mitochondrial dynamics are also altered in PAH as PASMCs isolated from PAH patients have increased activation of dynamin related protein-1 (Drp1) and decreased expression of Mfn2. Altered regulation of these proteins is associated with both increased mitochondria fragmentation and cell proliferation^7,31^. In Hela cells, overexpression of Sphk1 results in cleavage of Mfn2^7^. We observed that S1P leads to increased Drp1 activation and decreased Mfn2 expression, concomitant with increased PCNA expression, suggesting that the Sphk1/S1P signaling axis promotes PASMC proliferation via regulating mitochondrial dynamics.

In addition to regulating mitochondrial bioenergetics and dynamics, we observed that Sphk1/S1P signaling activates the mitochondrial stress response, the UPR^mt^. Sphk1 induced activation of the UPR^mt^ was first observed in *C. elegans* and the intestine, whereby Sphk1 was shown to bind to the outer mitochondrial membrane and activate ATF1 (ATF-5 is the mammalian homolog)^8^. We are reporting the novel observation that the effect of Sphk1/S1P on the UPR^mt^ is conserved across species. During homeostatic conditions, ATF-5 shuttles between the cytoplasm and mitochondria to perform mitochondrial surveillance. Decreased ATF-5 import into the mitochondria due to mitochondrial stress triggers nuclear translocation of ATF-5 and initiation of the UPR^mt^ transcriptional program (i.e. mtHSP70, HSP60 and ClpP) to restore mitochondrial homeostasis and promote cell survival^4^. Sphk1/S1P potentiated ATF-5 nuclear localization, as well as the expression and mitochondrial localization of ATF-5 target genes. The UPR^mt^ can be activated by any stimuli that triggers mitochondrial perturbations. Features that characterize the PAH microenvironment, such as hypoxia, oxidative-phosphorylation defects, and ROS, have been shown to promote the UPR^mt 31-37^. The mitochondrial toxin paraquat, leads to UPR^mt^ activation by interfering with electron transfer chain function at the inner mitochondrial membrane^9,31,38^. These reports combined with our observation that Sphk1/S1P promotes ox-phos defects, increases ROS generation and promotes ATF-5 nuclear translocation, suggests that the signaling axis promotes mitochondrial stress resulting in activation of the UPR^mt^.

ATF-5 also induces expression of genes that regulate the anti-apoptotic machinery, cell growth and migration ^4^. ATF-5 as well as its molecular targets, mtHSP70 and HSP60, have been linked to cell survival and growth in many types of cancer ^20,21,39-41^. Sphk1/S1P promoted PASMC proliferation, as we have previously reported. We used a specific, allosteric inhibitor of mtHSP70, Mkt-077, to inhibit UPR^mt^ signaling *in vitro* and *in vivo*. Inhibition of mtHSP70 reversed the Sphk1 mitigated increase in proliferation as we observed a decrease in PCNA expression. Furthermore, monitoring PASMC growth over a 48h time course demonstrated that Sphk1 overexpressing cells proliferate less in the presence of Mkt-077. We confirmed the effect of the UPR^mt^ on Sphk1 induced PASMC proliferation by knocking down expression of either ATF-5 or S1PR2. Deficiency in protein levels of both proteins significantly reduced proliferation. To evaluate the role of the UPR^mt^ *in vivo*, we first determined if the UPR^mt^ pathway is upregulated in HPH animal models. In PASMCS isolated from mouse lungs, we found increased expression of mtHSP70. In subsequent studies, we inhibited mtHSP70 in hypoxia exposed mice which reduced RVSP, RV hypertrophy and vascular remodeling compared to control mice.

Multiple types of mitochondrial dysfunction have been identified in PAH and have contributed to the metabolic theory of PAH pathogenesis. Herein, we report that the Sphk1/S1P/S1PR2 signaling axis contributes to vasculopathy that leads to a proliferative phenotype in PAH. These data demonstrate a profound pro-proliferative effect of Sphk1/S1P on mitochondrial function. This is the first report to demonstrate that this signaling axis activates the mitochondrial unfolded protein response and that targeting the UPR^mt^ mitigates HPH. We have shown that Sphk1/S1P individually regulates mitochondrial bioenergetics, dynamics, ROS and UPR^mt^. The UPR^mt^ can be activated by each of these mitochondrial pathways, however, further studies are necessary to investigate whether these pathways converge to promote vascular remodeling. These data suggest that targeting the UPR^mt^ may mitigate vascular remodeling in PAH but also reiterates the key role of mitochondria in the pathogenesis of PAH. Furthermore, they highlight the importance of elucidating the role of cooperative signaling among multiple mitochondrial processes and in understanding if pharmaceutically targeting one process affects other mitochondrial functions. This is a focus of our future studies.

## Figure Legends

**Supplementary Figure 1.**
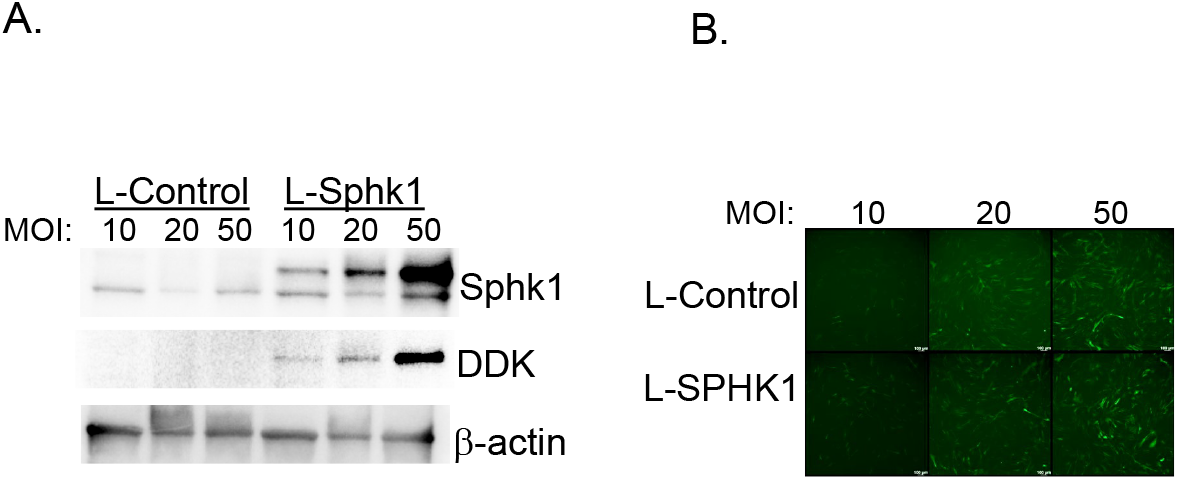
Overexpression of Sphk1 in hPASMCs. HPASMCs were transduced with Lenti-Sphk1 (MOI20, 48h). Western blot (A) and immunofluorescence for GFP (B) demonstrate lentiviral transduction and overexpression of DDK-tagged Sphk1. Results are expressed as mean ± SEM. ^*^p≤.05 versus time 0 or control.

